# A newly identified prophage-encoded gene, *ymfM*, causes SOS-inducible filamentation in *Escherichia coli*

**DOI:** 10.1101/2020.11.19.390815

**Authors:** Shirin Ansari, James C. Walsh, Amy L. Bottomley, Iain G. Duggin, Catherine Burke, Elizabeth J. Harry

## Abstract

Rod-shaped bacteria such as *Escherichia coli* can regulate cell division in response to stress, leading to filamentation, a process where cell growth and DNA replication continues in the absence of division, resulting in elongated cells. The classic example of stress is DNA damage which results in the activation of the SOS response. While the inhibition of cell division during SOS has traditionally been attributed to SulA in *E. coli*, a previous report suggests that the e14 prophage may also encode an SOS-inducible cell division inhibitor, previously named SfiC. However, the exact gene responsible for this division inhibition has remained unknown for over 35 years. A recent high-throughput over-expression screen in *E. coli* identified the e14 prophage gene, *ymfM*, as a potential cell division inhibitor. In this study, we show that the inducible expression of *ymfM* from a plasmid causes filamentation. We show that this expression of *ymfM* results in the inhibition of Z ring formation and is independent of the well characterised inhibitors of FtsZ ring assembly in *E. coli*, SulA, SlmA and MinC. We confirm that *ymfM* is the gene responsible for the SfiC^+^ phenotype as it contributes to the filamentation observed during the SOS response. This function is independent of SulA, highlighting that multiple division inhibition pathways exist during the stress-induced SOS response. Our data also highlight that our current understanding of cell division regulation during the SOS response is incomplete and raises many questions regarding how many inhibitors there actually are and their purpose for the survival of the organism.

**Importance:** Filamentation is an important biological mechanism which aids in the survival, pathogenesis and antibiotic resistance of bacteria within different environments, including pathogenic bacteria such as uropathogenic *Escherichia coli*. Here we have identified a bacteriophage-encoded cell division inhibitor which contributes to the filamentation that occurs during the SOS response. Our work highlights that there are multiple pathways that inhibit cell division during stress. Identifying and characterising these pathways is a critical step in understanding survival tactics of bacteria which become important when combating the development of bacterial resistance to antibiotics and their pathogenicity.

## Introduction

Bacterial cell division is an essential process that is tightly regulated to ensure division occurs at the correct time and position in order to create two viable and genetically identical daughter cells (1). In *Escherichia coli* this begins with the accumulation of the essential protein, FtsZ, into a ring-like structure (Z ring), at mid-cell (2). Following this, several downstream division proteins are recruited to form a complex, known as the divisome, which then constricts to divide the cell in two (3). There are several regulatory mechanisms that underlie the timing and positioning of division in *E. coli*. This includes the well-characterized Min system, which prevents the formation of Z-rings at the cell poles, and the nucleoid occlusion protein, SlmA, which inhibits Z-ring formation over unsegregated DNA (4, 5).

In addition to ensuring correct timing and positioning of the division site, there are numerous examples that demonstrate that the inhibition of division is equally important for cell survival under conditions such as DNA damage, protection from predation, progression of infection and pathogenesis (6–8). Inhibition of division results in the formation of filamentous cells; a process where cell growth and DNA replication continue in the absence of division, resulting in elongated cells (6). Filamentation is an important survival mechanism utilised by several bacteria in response to environmental stimuli (6, 8).

A well characterised cellular pathway that leads to filamentation is the SOS response which is activated by DNA damage under conditions including oxidative stress, antibiotic treatment or UV exposure (9). The activation of the SOS response is coordinated by two regulatory proteins, RecA and LexA (10, 11). RecA binds to ssDNA breaks caused by DNA damage, forming a RecA-DNA filament, which facilitates the self-cleavage of the LexA repressor, and the subsequent up-regulation (de-repression) of LexA-controlled genes (12). LexA represses over 40 genes in *E. coli* under normal growth conditions (13).

A key function of the SOS response is to inhibit cell division. This is thought to allow sufficient time for DNA repair to occur before committing to producing the next generation of daughter cells, minimising the transmission of defective DNA (14). In *E. coli*, this is facilitated by the cell division inhibitor, SulA, which is under the regulatory control of LexA (9). SulA is perhaps one of the most studied cell division inhibitors that causes filamentation, with molecular studies showing that it directly interacts with FtsZ, preventing assembly of the Z ring (15–19). When DNA damage is repaired, SulA is degraded via the cytoplasmic protease, Lon, and FtsZ polymerisation and cell division resumes (20, 21).

While SulA is generally the only cell division inhibitor commonly attributed to cell division inhibition during the SOS response in *E. coli*, other cell division inhibitors have been identified in this organism. Interestingly, several of these inhibitors include genes encoded within prophages (22–26). The e14 prophage contains an unidentified SOS-inducible cell division inhibitor, previously named SfiC+, which has been shown to contribute to filamentation during the SOS response, independent of SulA (27, 28). This inhibitor was not however under LexA repression (27). Phages contain their own repressor systems, which are LexA-like in nature, such as the CI-repressor from λ phage (13, 29). The gene *cohE* from e14, encodes a CI-like repressor, similar in sequence to other bacteriophage CI repressors that are responsive to an SOS signal (30). As such, it is possible that expression of *sfiC* is regulated by *cohE*. In earlier work, it was also shown that FtsZ was most likely the target of SfiC+, as point mutations in *ftsZ* that confer resistance to the inhibitory effects of SulA, also conferred resistance to SfiC+ (27, 28). Based on prophage arrangement, the gene responsible for the SfiC+ phenotype has been suggested to be likely encoded by either of the adjacent e14 prophage genes *ymfL* or *ymfM* (30), however the precise identity of the gene remains unknown.

We previously developed a high-throughput flow cytometry system to screen a novel *E. coli* expression library for candidate cell division regulators (31). We identified a DNA fragment containing the e14 prophage-encoded genes, the full *ymfM* gene and a partial sequence of adjacent genes, *ymfL* and *oweE* (previously annotated as *ymfN*), that when expressed in an inducible plasmid-based system, caused cells to elongate and form filaments (31, 32). Since this DNA fragment contained both potential candidates for SfiC+, *ymfM and ymfL*, further work was needed to determine which of these genes is responsible for the SfiC+ phenotype i.e. SulA-independent filamentation during the SOS response.

Here we show that *ymfM* is responsible for the SfiC phenotype, resulting in the inhibition of cell division when expressed from an inducible plasmid. We further characterise the effect of *ymfM* expression on *E. coli* cell division and show that results in prevention of the early stage of cell division: Z-ring formation. Furthermore, its inhibition pathway is independent of known cell division inhibitors, SulA, SlmA and MinC. Finally, we show that YmfM causes filamentation during activation of the SOS response and that this is independent of SulA. Our data indicate that multiple division inhibitors exist during the SOS response and raises questions regarding how their purpose for the survival of the organism.

## Results

### 1. Identification of a new gene, *ymfM*, whose expression induces filamentation

Previous independent theoretical (30) and experimental (27, 31) studies have identified that either *ymfM* or *ymfL*, is *sfiC*. To determine which one of these genes is responsible for the SfiC+ filamentous phenotype, *ymfM* and *ymfL* were cloned separately into the arabinose-inducible plasmid, pBAD24. Cells were grown in LB supplemented with 0.2% glucose to mid-exponential phase to repress gene expression. The cells were then diluted in fresh LB supplemented with 0.2% arabinose to induce gene expression and then grown for at least 4 doubling times (generation time is approximately 30 minutes) to allow for enough time to observe cell length changes. The degree of filamentation associated with the expression of each individual gene was measured as cell length (μm) from phase-contrast microscopy images. Control cells expressing just pBAD24 had an average cell length of 4.2 ± 1.5 μm, and these ranged from approximately 2 μm to 10 μm in length (Figure S1). Therefore, filamentous cells resulting from gene expression in pBAD24 were defined as being greater than 10 μm in cell length.

The induction of *ymfL* expression did not cause filamentation, with cells having an average length of 3.5 ± 0.9 μm (Figure 1A). However, expression of *ymfM* resulted in inhibition of division, giving rise to an exclusively filamentous population having an average cell length of 57.3 ± 19.7 μm (Figure 1B). In the repressed state, cells had the same cell-length distribution as the empty vector (Figure 1 and S1). We propose that cell division is stopped abruptly in *ymfM*-induced cells. The initial doubling time was 30 min, which for the 120-min total incubation would result in filamentous cells 16-fold (2^4^) longer than their short-cell counterpart. However, we noticed that after 90 min of the 120-min incubation period that *ymfM*-induced cells entered stationary phase (data not shown). This resulted in a mean length of *ymfM*-induced cells of 14.5 times the *ymfM*-repressed mean of 3.9 μm. Overall these results identify conclusively that *ymfM* is the gene responsible for the filamentation observed in the *E. coli* overexpression screen (31) and in the SfiC+ phenotype (27). They also indicate that this induced expression of *ymfM* results in an earlier onset of stationary phase compared to repressed conditions.

**Figure 1.**
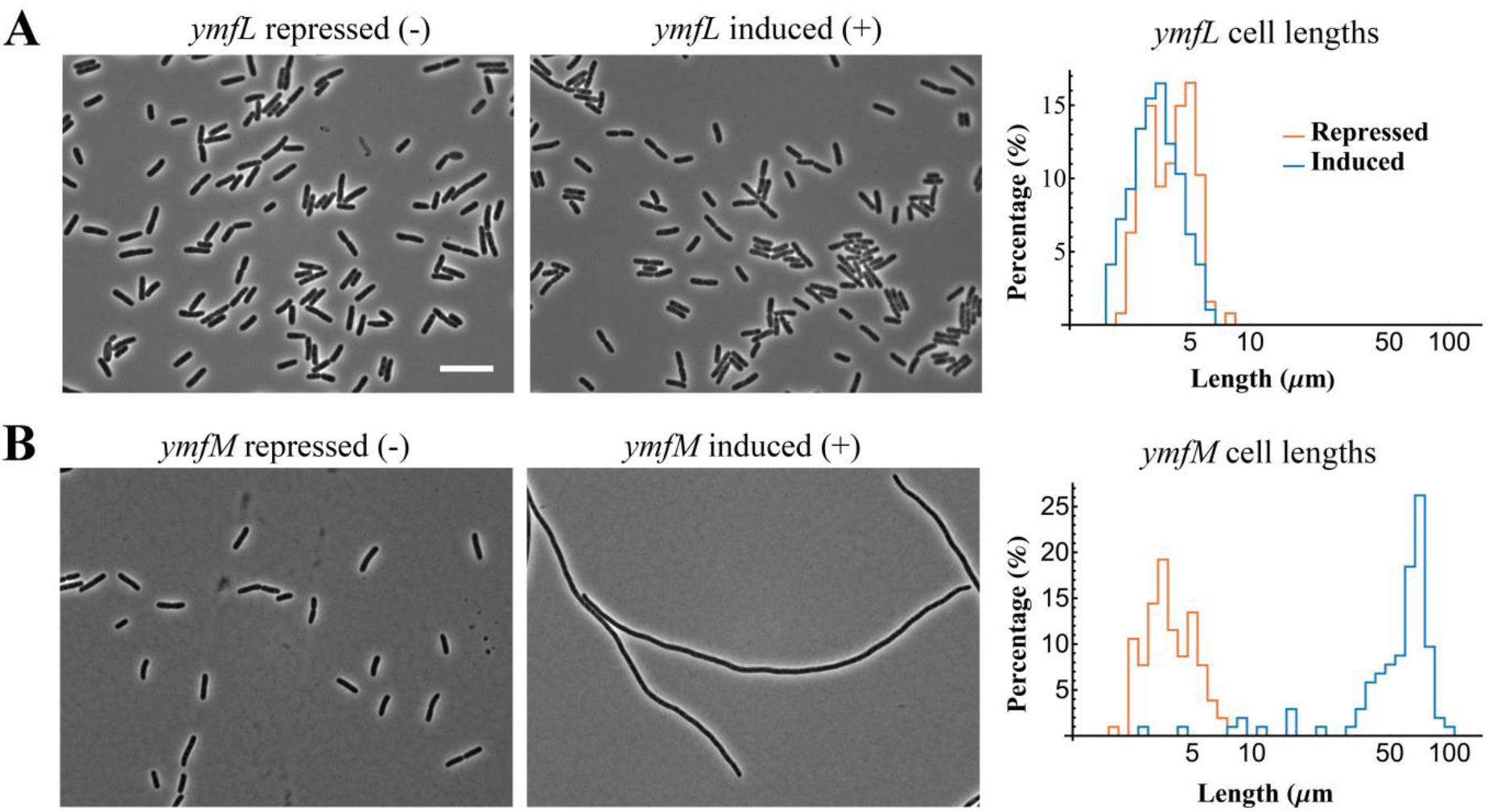
Cell length distribution when *ymfM* or *ymfL* is expressed from an inducibleplasmid. The open reading frame for genes (A) *ymfL* and (B) *ymfM* were cloned into the arabinose-inducible plasmid, pBAD24, and grown in LB medium, supplemented with 0.2% (w/v) glucose to repress their expression or 0.2% (w/v) arabinose to induce expression, for two hours (4 generations). Representative phase-contrast image of induced *ymfM* expression shows filamentation of cells while the expression of *ymfL* does not affect cell length. Approximately 100 cells were measured for each population and their respective cell lengths overlayed in a histogram shown on the right. Scale bar = 10μm

### 2. *ymfM* expression inhibits the earliest stage of division: FtsZ ring formation

To understand whether induction of *ymfM* expression inhibits cell division by impeding FtsZ assembly or a later stage of cell division, immunofluorescence microscopy (IFM) was used to measure Z ring frequency in cells induced to express *ymfM* from pBAD24. The absence of Z rings in the *ymfM*-induced cells would imply that Z ring formation has been inhibited.

IFM using cephalexin-treated filamentous cells was first performed as a control to show that the technique did not affect the integrity of the Z rings in filaments, and other control experiments showed that the antibody detection for IFM is specific for FtsZ (Figure S2).

Next, wild-type cells harbouring pBAD-*ymfM* were grown in LB for 3 generations with either glucose to repress expression or with arabinose to induce expression for 90 min. During repressed conditions, cells had an average cell length of 3.5 ± 0.9 μm indicating that they were dividing at the normal frequency. In these short cells, Z rings were observed as bright green bands at mid-cell (white arrow, Figure 2), at a frequency of 8.4 μm of cell length per Z ring. In cells expressing *ymfM*, the average cell length was 35.5 ± 23.1 μm and almost no Z rings were observed along the length of the filament. Instead FtsZ appeared to be diffused throughout the filament (Figure 2B). Occasional Z rings were seen in this sample; however these were primarily in the few short cells present. Z rings in this population were observed at a frequency of 224 μm of cell length per Z ring – 27-fold less frequent compared to repressed cells. DAPI (DNA) staining showed that in *ymfM*-induced filaments, that the nucleoids appeared normal, suggesting that the filamentation observed is solely due to the inhibition of division and not a result of DNA replication or chromosome segregation inhibition. Furthermore, it was shown that the lack of Z-rings in filaments was not due to changes in cellular levels of FtsZ (Figure S3).

**Figure 2.**
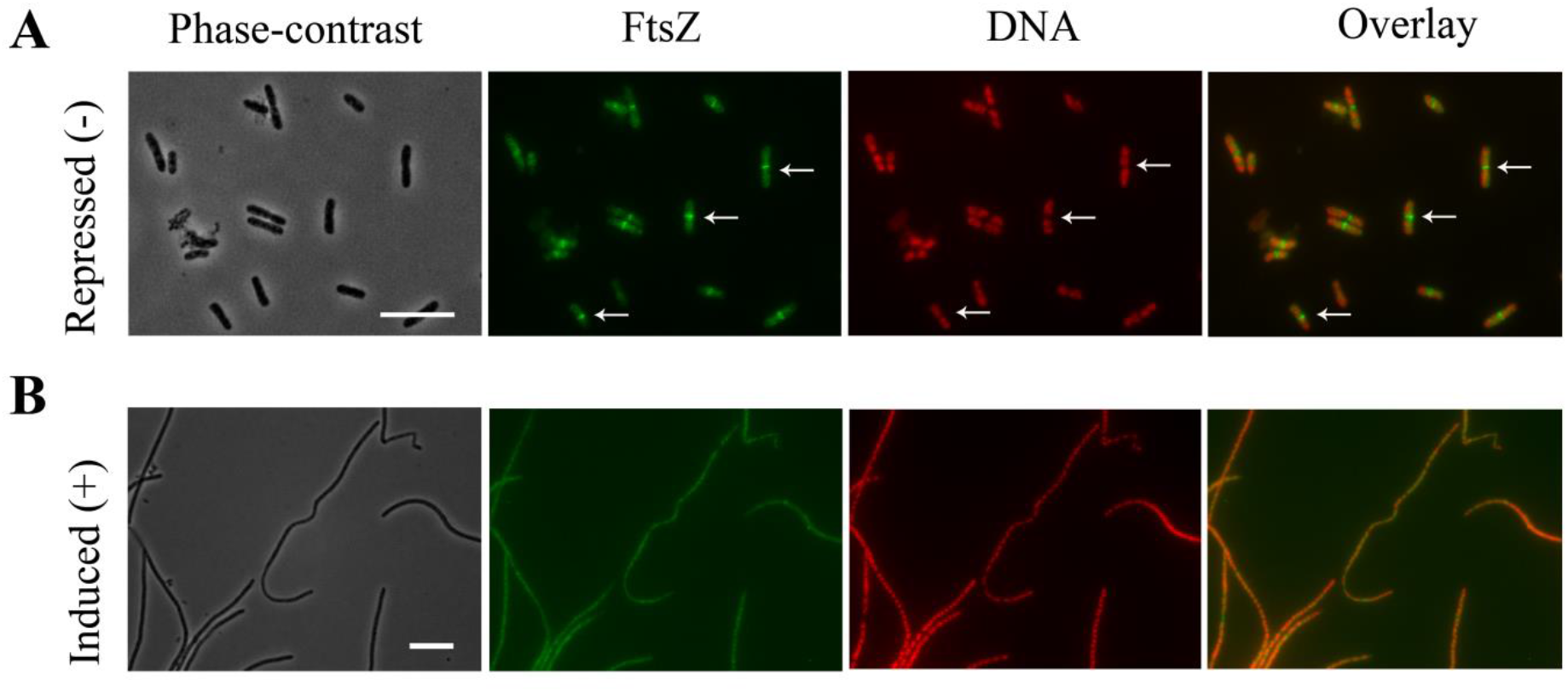
Z ring assembly is inhibited in filamentous cells induced by *ymfM* expression. Immunofluorescence microscopy using anti-FtsZ antiserum to visualize Z rings of strain BW25113 carrying pBAD-*ymfM*. When grown in LB supplemented with 0.2% (w/v) glucose to repress *ymfM* expression (A) short cells are present with Z rings at mid-cell (white arrows). Expression of *ymfM* was induced with 0.2% (w/v) arabinose for three generations (90 min) and the resulting filamentous cells contain no Z rings (B). The nucleoids have been stained with DAPI (falsely coloured red) and the overlay image shows Z-ring positioning within cells relative to nucleoids. Scale bars = 10μm.

In summary, induced expression of *ymfM* results in inhibition of Z ring formation at midcell and this is not due to change in the cellular levels of FtsZ.

### 3. YmfM inhibition of division does not rely on known cell division regulators, SulA, SlmA and MinC

Induced expression of *ymfM* inhibits the earliest stage of division as Z rings were not detected with IFM (Figure 2). We next tested whether this inhibition occurred through other Z ring regulators. These include SulA, which inhibits FtsZ from assembling into a ring during the SOS response (15, 16); MinC, which prevents FtsZ assembly at cell poles under normal conditions (33, 34); and SlmA, which prevents FtsZ assembly over nucleoids as part of the nucleoid occlusion system in *E. coli* (35, 36). The expression of *ymfM* was induced in *E. coli* cells in the absence of SulA, (Δ*sulA*; JW0941) (37), SlmA (Δ*slmA*; JW5641) (37) or the Min system (Δ*minCDE*; TB43) (35). If YmfM prevents Z ring assembly specifically via any one of these other inhibitors, then no filamentation will be observed in their absence when *ymfM* is expressed.

When *ymfM* expression was repressed, the mutant strains had a short cell distribution which was comparable to their wild-type counterpart, except for Δ*minCDE*. The absence of the Min proteins is reported to cause a mixed population of minicells (small cells lacking DNA) and mildly filamentous cells due to polar cell division (15) and this was observed in the Δ*minCDE* cell population. Cell lengths were 3.9 ± 1.2 μm (BW25113, parent of Δ*sulA* and Δ*slmA*), 3.8 ± 1.0 μm (Δ*sulA*), 5.7 ± 4.2 μm (Δ*slmA*), 4.3 ± 1.4 μm (TB28, parent to Δ*minCDE*) and 10.6 ± 5.5 μm (Δ*minCDE*).

When *ymfM* expression was induced in the mutant strains, Δ*sulA*, Δ*slmA* or Δ*minCDE*, filamentation was observed (Figure 3A). The average cell lengths were of 71.8 ± 19.7 μm (Δ*sulA*), 70.6 ± 28.3 μm (Δ*slmA*) and 87.7 ±16.1 μm (Δ*minCDE*), respectively. Filamentation in the mutant strains was comparable to their respective wild-type parent strain (Figure 3B), with average cell lengths of 76.2 ± 10.8 μm for BW25113 and 93.9 ± 15.4 μm for TB28. The results here show that *ymfM* does not require the inhibitory activity of SulA, SlmA or the Min system to inhibit Z ring formation.

**Figure 3.**
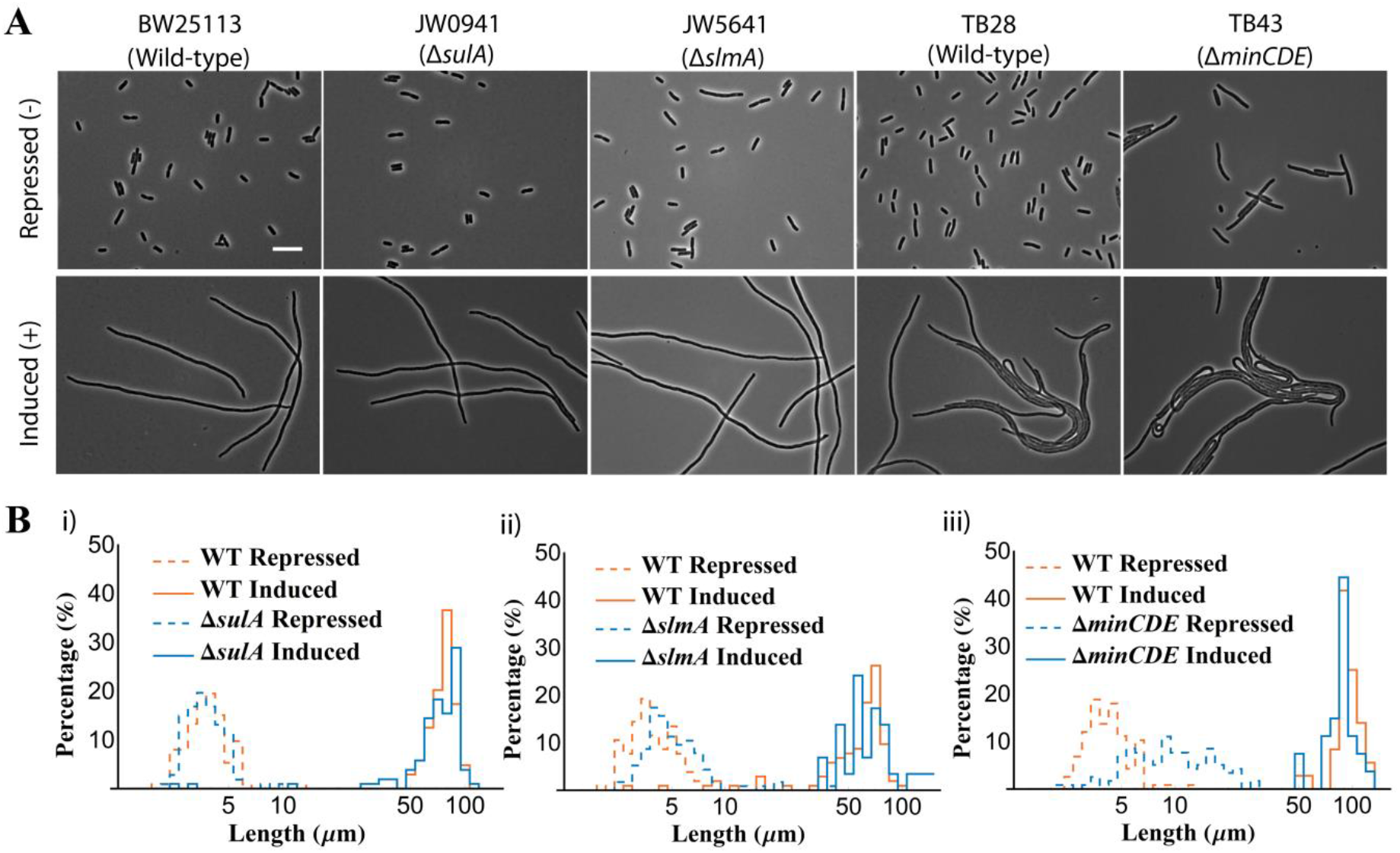
Filamentation caused by the expression of *ymfM* is independent of the cell division inhibitors, SulA, SlmA and the Min system. A) Phase-contrast images of strains, Δ*sulA* (JW0941), Δ*slmA* (JW5641), Δ*minCDE* (TB43) and their wild-type backgrounds (BW25113 and TB28, respectively) show filamentation when *ymfM* expression is induced from pBAD24 with 0.2% (w/v) arabinose in LB for two hours (4 generations). Short cells are observed when *ymfM* expression is repressed with 0.2% (w/v) glucose, with the exception of Δ*minCDE*, which has a mixed population of short, slightly filamentous and minicells due to increased division at cell poles. B) Cell length histograms of the mutants, (i) Δ*sulA*, (ii) Δ*slmA* and (iii) Δ*minCDE* show that the degree filamentation caused by *ymfM* expression is comparable to their wild-type counterparts. Approximately 100 cells were counted for each cell population. Scale bar = 10μm

### 4. YmfM is involved in the inhibition of division during the SOS response and is independent of SulA

Several years ago, an unknown gene, *sfiC*, was identified as a cell division inhibitor during the SOS response (27). D’ Ari *et al*. (27) found that even in the absence of the *sulA* gene, cells were able to filament when SOS was activated as *sfiC* would inhibit cell division. We have thus far shown that *ymfM* is *sfiC*, as its expression from an inducible plasmid causes filamentation (Figure 1) and this is independent of the inhibitory actions of SulA (Figure 3). *ymfM* should therefore also be responsible for the SulA-independent filamentation observed during the SOS response.

To show *ymfM* was required for filamentation during SOS in the absence of *sulA*, we used a similar approach to D’Ari et al (27), in which *sfiC* was identified using a temperature-sensitive mutant, *recA441*, also known as *recA-tif* (38). In this mutant, RecA acquires protease activity at the non-permissive temperature of 42°C, thereby inducing the SOS response and filamentation without the need for external means to DNA damage (39). We therefore cloned *recA441* into pBAD24. Gene expression and subsequent SOS was induced with 0.2% arabinose and growth at 42°C in strains Δ*ymfM*, Δ*sulA*, or Δ*sulA*Δ*ymfM*. Any changes to the degree of filamentation were measured. If, in the absence of *ymfM*, cells do not filament as effectively as their wild-type counterparts, this would suggest that *ymfM* directly contributes to filamentation during the SOS response.

As expected, induction of *recA441* caused filamentation in the wild-type background (Figure 4, yellow line) as compared to the short cell population of wild-type cells expressing empty vector (blue line). Here, cell size (reported as μm^3^) was measured using a Coulter cytometer and this is proportional to cell length as cell width is unchanged.

**Figure 4.**
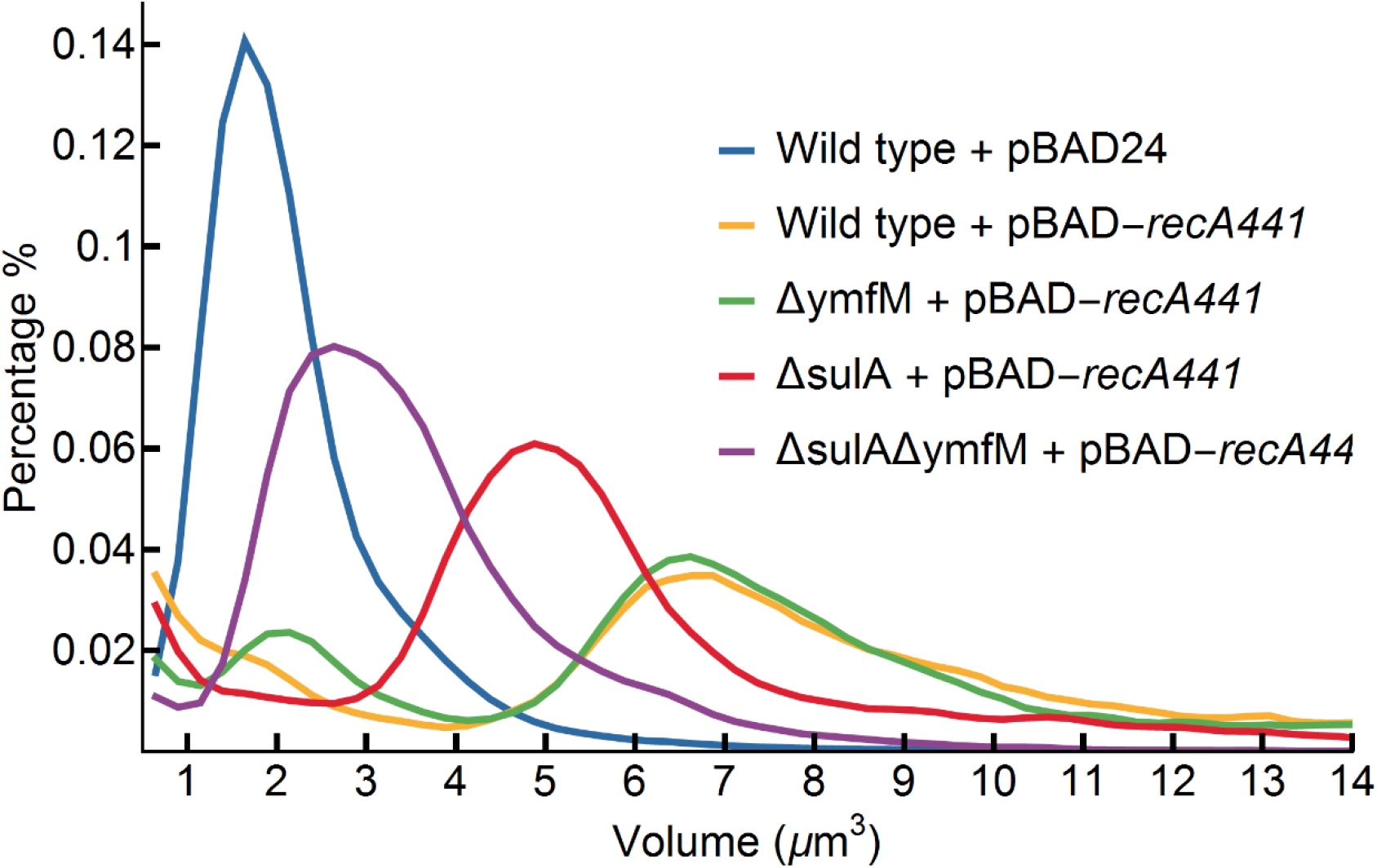
YmfM contributes to the filamentation of *E. coli* during induction of the SOS response using the temperature-sensitive RecA mutant, *recA441*. Coulter cytometer analysis of cell size distribution of wild-type (yellow), Δ*ymfM* (green), Δ*sulA* (red), and Δ*sulA*Δ*ymfM* (purple) after two hours of pBAD-*recA441* induction in LB with 0.2% (w/v) arabinose and 100 μg/mL adenine, at 42°C. Samples were compared to wild-type cells expressing empty pBAD24 (blue) not under SOS induction. X-axis represents cell volume (μm^3^) and Y-axis represents the percentage of cells at any given volume. Pulse data > 10,000 events (cells). Data represents one biological replicate. Three additional biological repeats were performed (Fig S4).

There was a reduction in the degree of filamentation in the absence of *sulA* alone (Figure 4, red line), as compared to its wild-type counterpart. However, this did not fully recover to the short cell population (blue line), highlighting the presence of additional cell division inhibitor(s) that are active during the SOS response. The absence of *ymfM* (green line) was similar to that of wild-type (yellow line), so the absence of *ymfM* alone does not result in less filamentation in the population. However, in the absence of both *sulA* and *ymfM* (purple line), there was the greatest shift towards the shorter cell population (blue line). The difference in cell size distribution of Δ*sulA*Δ*ymfM* (purple line) cells compared to Δ*sulA* (red line) cells shows that *ymfM* does indeed play a role in inhibiting division during the SOS response when *recA441* is induced and this role becomes evident when SulA is absent. Importantly, while the shift towards shorter cells was the greatest for Δ*sulA*Δ*ymfM* (purple line), it did not lead to a full recovery to a short cell population (blue line), suggesting the presence of additional SOS-inducible inhibitors in this organism. These trends were observed in multiple biological replicates (Figure S4).

## Discussion

We have identified the gene, *ymfM*, which when expressed from an inducible plasmid causes arrest in cell division (Figure 1). YmfM is encoded within the e14 prophage which had been indicated to harbour the gene responsible for the SOS-inducible filamentation phenotype, SfiC+ (27, 28). The exact gene responsible for SfiC+has remained elusive for over 35 years and was narrowed down to either *ymfL* or *ymfM* in 2004 (30). In this work, we eliminate *ymfL* and confirm *ymfM* as being responsible for the inhibition of division, confirming that *ymfM* is responsible for the filamentous SfiC+ phenotype reported in 1983 (27).

We first characterised the stage of division being inhibited by *ymfM*. The expression of *ymfM* inhibits Z ring formation as essentially no rings were seen in cells expressing *ymfM*. This is consistent with previous work by D’ Ari *et al*., (27) who demonstrated that mutations in *ftsZ* that confer resistance to the inhibitory effects of SulA also confer resistance to SfiC. We also show that YmfM does not act through known *E. coli* cell division regulators, SulA, SlmA or MinC. Another phage-encoded inhibitor, DicB from prophage Qin has been reported to utilise the inhibitory actions of MinC as no filamentation by DicB was observed in the absence of the Min System (40, 41). Filamentation by YmfM was observed in the absence of *sulA, slmA* and *minCDE*, showing that it is independent of these inhibitory pathways. It remains to be seen whether *ymfM* targets FtsZ directly to inhibit Z ring assembly, or it does so through early division proteins such as FtsA or ZipA (3), similar to the phage inhibitor, Kil (22, 42). Our data do not rule out the possibility that YmfM causes division inhibition indirectly through an as yet unidentified cell division inhibitor.

Since we have shown that induction of *ymfM* expression causes filamentation and is responsible for the SfiC^+^ phenotype, then, as an SOS-inducible gene, it should also contribute to filamentation when the SOS response is induced through the activation of RecA, as reported by D’ Ari *et al* (27). This indeed was the case. Through activation of *recA441*,YmfM inhibited division during the SOS response (Figure 4). This was also apparent when the degree of filamentation is compared between Δ*sulA* and Δ*sulA*Δ*ymfM*, as the double-knockout resulted in shorter filaments as compared to Δ*sulA* alone. This difference in filamentation can be attributed to *ymfM*.

The absence of *sulA* alone was enough to show a partial reduction in filament size, suggesting that SulA is the primary inhibitor of cell division during the SOS response. The lack of change in cell size in Δ*ymfM* as compared to wild-type was unexpected given that expression from *pBAD-ymfM* causes very filamentous cells. This may be due to differences in expression levels of *ymfM* from the plasmid versus through SOS induction. Furthermore, it is likely that in the SOS-induced system, as SulA is still present, SulA and potential additional cell division inhibitors are masking the phenotypic effects that can be caused by the absence of *ymfM*.

It was also apparent that genes additional to *sulA* and *ymfM* contribute to filamentation during RecA-activated SOS, as there was not a full recovery to a short-cell population in the Δ*sulA*Δ*ymfM* background (Figure 4). Given that over 1000 genes are differentially expressed during the SOS response (43), it is likely that several SOS-inducible cell division inhibitors are yet to be identified. For example, KilR, another prophage-encoded cell division inhibitor, has only recently been shown to be activated by the small RNA, *oxyS*, in response to oxidative stress (44).

It is interesting to speculate why multiple cell division inhibitors are present during the SOS response and ask, how may they differ to SulA? Additionally, why are so many of these division inhibitors present in prophages? It has been shown that the 9 cryptic prophages present in *E. coli* K-12 are beneficial for survival and adaption under different environmental conditions and signals, including osmotic, oxidative and acid stress, biofilm formation and tolerance to antibiotics (28, 30, 45). Several of these prophages also encode cell division inhibitors such as Kil, DicB, DicF, and CbtA (23–26). *KilR* from prophage Rac and *dicB* from prophage Qin are both cell division inhibitors which have been shown to be up-regulated during treatment with nalidixic acid and azlocillin and are thought to contribute to the resistance of these antibiotics (24, 25, 45). It is possible that these prophage-encoded inhibitors, originally serving the functions of the phage, have more recently been adapted to be induced in response to specific environmental cues for the benefit of the bacterium (45, 46).

It was of interest to us to understand conditions under which *ymfM* is active as this may help us differentiate how its function differs to that of SulA. Expression of *ymfM* is up-regulated during norfloxacin induced SOS (47–49). However, while we could also show that *ymfM* expression was upregulated under these conditions, we were repeatedly unable to show the requirement for this gene in causing filamentation in the presence of this antibiotic (data not shown) as we observed in the RecA441 experiments (Figure 4). As with the *recA441* experiments, we think it likely that the *ymfM* phenotype in the presence of norfloxacin is being masked by numerous filamentation mechanisms. Alternatively, since *ymfM* is not under the control of LexA repression (27), and likely controlled by the CI-like repressor within the e14 prophage, it possible that *ymfM* responds to different yet-to-be identified conditions. These observations also highlight the possibility that multiple cell division inhibitor genes and pathways exist for a more finely tuned division regulation response to the environment. For example, spatial inhibitors MinC and SlmA have specialised functions to ensure division does not occur at inappropriate times in the cell cycle or the incorrect position in the cell. It is probable that temporal inhibitors are equally specialised with respect to when they are activated. By having multiple inhibitors with subtly different properties, cells are likely able to tailor their inhibition in response to different environmental conditions and signals to maximise their survivability. More work is needed to tease apart the relationship between different inhibitors and the conditions under which they are required.

Overall, our data shows that *ymfM* is a novel gene required for cell division inhibition during the SOS response, and its activity is independent of a major SOS-induced cell division inhibitor, SulA. Our data also highlights that our current understanding of the cell division regulation during the SOS response is incomplete and raises many questions. In particular, what are the benefits of having multiple cell division inhibitors during the SOS response? How are the different inhibitors activated? And how do these multiple pathways help *E. coli* cope with stresses and aid in the survival of the population, if at all?

## Methods and Materials

### Strains and Growth Conditions

All *E. coli* strains used in this study are listed in Table S1. *E. coli* cells were grown in LB media with vigorous shaking (250 rpm) at 37°C, unless stated otherwise. Ampicillin (100μg/mL; Sigma Aldrich) was supplemented, where appropriate. Lambda Red recombination (37, 50) was used to generate gene deletions in *E. coli* strains listed in Table S1.

### Plasmid construction and expression

All plasmids used in this study are listed in Table S2. Recombinant plasmids were constructed using the Gibson assembly method (51) following the manufacturer’s instructions for the Gibson assembly master mix (NEB). DNA fragments *ymfM, ymfL* and *oweE* were amplified from BW25113 genomic DNA and *recA441* was amplified from JM12 genomic DNA (38), see Table S3. Each DNA fragment contained homologous overlap to pBAD24 which was linearised at the *NcoI* restriction site.

### Expression of DNA fragments from pBAD24

Cultures of desired *E. coli* strain containing recombinant pBAD24 were grown overnight in 5mL LB with ampicillin (100 μg/mL) and 0.2% glucose (repression of araBAD promoter) at 37°C, or 30°C for pBAD-*recA441*. Cultures were diluted to OD_600_= 0.04 in 20mL LB with ampicillin (100 μg/mL) and 0.2% glucose and grown at 37°C (or 30°C for pBAD-*recA441*), 250 rpm to mid-exponential phase (OD600 ~ 0.5). A 1mL aliquot of the culture was collected and fixed with 3% formaldehyde. Remainder of the culture was pelleted by centrifugation at 2, 000 x *g*, and washed twice in an equal volume of fresh LB to remove the glucose. Cultures were further diluted to an OD_600_=0.04 in 20mL LB with ampicillin (100 μg/mL) and 0.2% arabinose (to induce expression) and grown for 4 generations (approximately 2 hours) at 37°C with shaking. For *recA441* expression, cultures were grown at 42°C and supplemented with 100 μg/mL adenine in addition to the arabinose and ampicillin.

### Microscopy

#### Immunofluorescence microscopy (IFM)

IFM was used for the detection of the FtsZ protein and is based on the method previously described in (52), with the exception that cell lysis was omitted. Cultures of BW25113 containing pBAD-*ymfM* were grown as detailed above and incubated with the primary antibody, αFtsZ (anti-sera), diluted 1:10,000 in BSA-PBS, at 4°C overnight. The primary antibody was removed, and the cells were incubated with Alexa488-conjugated secondary antibody, αRabbit IgG (Invitrogen), diluted 1: 10,000 in BSA-PBS, for 2 hours in the dark, at room temperature. Wells were washed with PBS to remove excess secondary antibody and DAPI (4’6-diamidino-2-phenylindole), at a final concentration of 2μg/mL was added to each sample. Cell morphology and FtsZ localisation was then examined using phase-contrast and fluorescence microscopy as described below.

#### Phase-contrast and Fluorescence Microscopy

Cells were imaged using phase-contrast and fluorescence on a Zeiss Axioplan 2 fluorescent microscope equipped with a Plan Apochromat (100x NA 1.4; Zeiss) objective lens. The light source was a 100 W high pressure mercury lamp passed through the following filter blocks for visualising Alexa Fluor 488 (Filter set 09, Zeiss; 450 – 490 nm BP excitation filter, 515 nm long pass (LP) barrier filter), and for visualising DAPI (Filter set 02, Zeiss; 365 nm excitation filter, 420 nm long pass (LP) barrier filter). Images were collected using the AxioCamMRm camera and processed using the AxioVision 4.8 software (Zeiss). Approximately 100 cells were measured from each data set (unless specified otherwise) using the length tool within the Axiovision software.

#### Coulter cytometer

A volume of 100μL of fixed cells was added to 9.9mLs of Isoflow buffer (Beckman). Of this, 200μL was run through a 50μm aperture tube, and data collected over 400 bins ranging from 0.6 μm^3^ to 100μm^3^. Data was exported in excel and plotted as a histogram with the cell volume along the x-axis and percentage of cells on the y-axis, in Mathematica (Wolfram).

## Acknowledgements

This work was not specifically funded by a research grant. The funding to support this work was provided by the institution of the corresponding author (University of Technology, Sydney, Australia). S.A. was supported by an Australian Government Research Training Program Stipend. We thank Tom Bernhardt for strains *E. coli* strains TB28 and TB43, Iain Duggin for strain JM12 and plasmid pBAD24, Joe Lutkenhaus for strain JKD-2, and Piet de Boer for advice on immunofluorescence protocols with *E. coli*.

